# Noise-induced properties of active dendrites

**DOI:** 10.1101/2020.11.30.405001

**Authors:** Carl van Vreeswijk, Farzada Farkhooi

**Affiliations:** Centre de Neurophysique Physiologie et Pathologie, Paris Descartes University and CNRS UMR 8002 INCC, 75006 Paris, France; Institute for Theoretical Biology, Department of Biology, Humboldt-Universität zu Berlin, 10115 Berlin, Germany

**Keywords:** Noise-induced bistability, Noise-induced non-monotonicity, Neuronal dendrites, Voltage-gated calcium channels, Slow-fast analysis, Fokker-Planck equation

## Abstract

Dendrites play an essential role in the integration of highly fluctuating input into neurons across all nervous systems. Nevertheless, they are often studied under the conditions where inputs to dendrites are sparse. Up to date, the dynamic properties of active dendrites facing in-vivo-like fluctuating input remains elusive. In this paper, we uncover fundamentally new dynamics in a canonical model of a dendritic compartment with active calcium channels, receiving in-vivo-like fluctuating input. We show in-vivo-like noise induces non-monotonic or bistable dynamics in the input-output relation of a dendritic compartment, both of which are absent in a noiseless condition. Our analysis shows that the timescales of the activation gating variable of the dendritic calcium dynamics determine noise-induced spontaneous order in the system. Noise can induce non-monotonicity or bistability with fast or slow calcium activation respectively. We characterize these noise-induced phenomena and their influence on the input-output relation. Furthermore, we show that timescales of the emerging stochastic bistable dynamics go far beyond a deterministic system due to stochastic switching between the solutions. Our results reveal that noise contributes to sustained dendritic nonlinearities, and it could be considered a principal component of the dendritic input integration strategies.

---

The interactions among neurons are mediated via synapses primarily located on large tree-like dendritic structures. Dendritic trees integrate these synaptic inputs and determine the extent to which the neuron produces spiking output [1]. Dendrites contribute substantially to neuronal plasticity and functions [2–4], and their active calcium dynamics can induce nonlinear regenerative events such as dendritic spikes in-vitro [5] and in-vivo [6] conditions. The dendritic spikes have also been shown to serve a functional role in in-vivo cortical visual processing, where a typical pyramidal neuron receives intense fluctuating input, generated by the summation of hundreds to thousand synaptic inputs from excitatory and inhibitory presynaptic neurons in the circuit [7]. Although the dendrites’ importance for integration of the relevant signal in-vivo conditions is evident [3], how a dendritic compartment shapes its input-output relation in this noisy condition is essentially unknown. While noise is commonly assumed to be a nuisance that has to be filtered out, it can also induce new organized behaviors in systems, which are absent in deterministic conditions [8]. Examples where noise leads to the emergence of spontaneous order in biological and physical systems include stochastic resonance [9], noise-induced phase transitions [10], and noise-induced bistability [11]. To investigate whether in-vivo-like fluctuating input also induces novel dynamic states in dendrites, we analytically study a canonical model of a dendritic compartment with active calcium channels. Our results indicate that stochastic phenomena emerge in dendritic dynamics with in-vivo-like fluctuating input, which is entirely absent in the deterministic condition. We uncover that the noise induces non-monotonic or bistable dynamics in the dendritic input-output relation, depending on the timescales of calcium-gating variables.

## The dendritic compartment model

To study the noise-induced dynamics in dendrites, we consider the canonical conductance-based description of the dendritic compartment membrane potential voltage, *v*, as expressed by

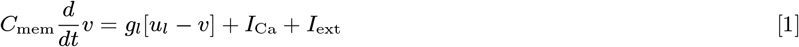

where, *C*_mem_ and *g_l_* are the membrane capacitance and leak conductance per unit area, *u_l_* is the leak reverse potential, and *I*_Ca_ is the active dendritic feedback. We model in-vivo-like external input per unit area, *I*_ext_, by a generic white-noise process as

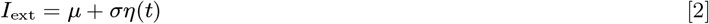

where, *μ* and *σ* are the mean and standard deviation of the process, and *η*(*t*) is a Gaussian white noise variable, where 〈*η*(*t*)〉 = 0 and 〈*η*(*t*)*η*(*t′*)〉 = *δ*(*t* − *t′*). We use the notation of 〈.〉 to denote averaging over the noise realization. The dendritic active feedback properties due to calcium dynamics in Eq. (20) is captured by

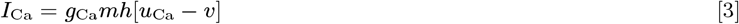

where, *g*_Ca_ and *u*_Ca_ are the maximum calcium conductance per unit area and its reverse potential, respectively. The dynamics of the activation gating variable, *m*, and inactivation gating variable, *h*, are given by and

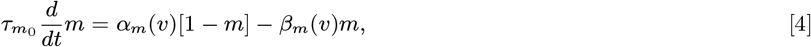

and

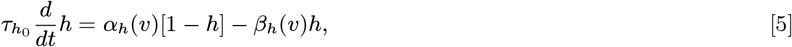

respectively. In Eq. (23) and Eq. (24), the channel opening occurs at a voltage-dependent rate *α_x_*(*v*) and the closing rate is *β_x_*(*v*), where *x* ∈ {*m, h*}; and the thermodynamic model for voltage-dependent currents [12] suggests 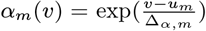, 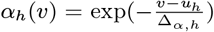, 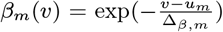 and 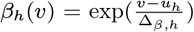, where *u_x_* is half-activation or -inactivation potential, and Δ*_α,x_* and Δ*_β,x_* are slope factors for opening and closing of the channel. To reduce the number of parameters, we assume Δ*_α,m_* = Δ*_α,h_* = Δ*_α_* and Δ*_β,m_* = Δ*_β,h_* = Δ*_β_*. The time-constant of the activation and inactivation gating variables are tuned by 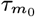 and 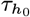, respectively. For numerical analysis and simulations, the model parameters’ values are given in Table 1, unless it is otherwise specified.

**Table 1.**
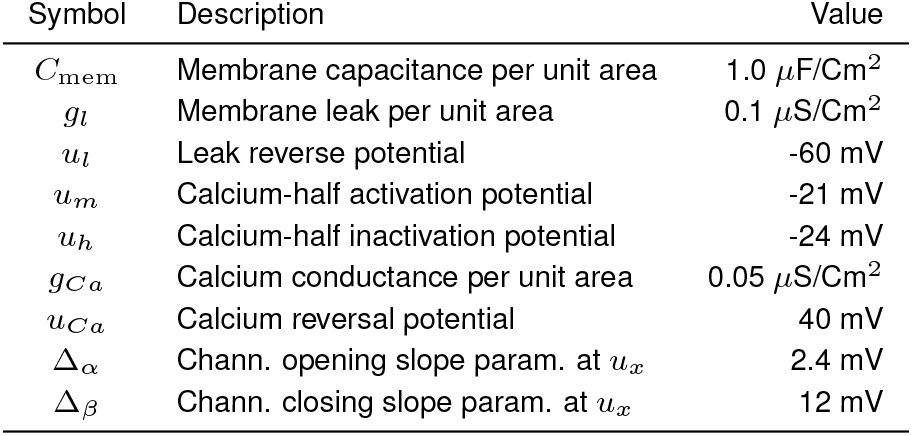
The dendritic compartment model parameters

In a noiseless dendritic compartment with a constant input *μ*, the equilibrium dendritic voltage depends on steady-state values of *m* and *h* which are denoted by 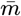 and 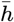, and it is given by 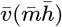, where

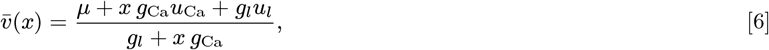

and self-consistency requires 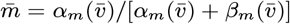 and 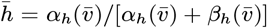. In Figure 1, we plot the voltage equilibrium value, 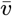, as the function of constant input *μ*. The 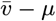 function exhibits a monotonic increase in the average dendritic voltage 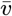, when *μ* increases with small acceleration when calcium feedback is sufficiently activated. Figure 1 conveys that the equilibrium value of dendritic voltage in the deterministic system does not depend on the effective timescales of the calcium gating variables, given by 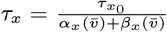 (note that *x* ∈ {*m, h*}). The dendritic calcium dynamics are known to be much slower than the membrane time-constant, *τ*_mem_ = *C*_mem_*/g_l_* [13]. In particular, the time-constant of the inactivation variable, *h*, is large compared to the membrane [13] and the dynamics of the activation variable *m* is small for some calcium channels [13], while it is large for others [13–15]. In this paper, we show that these timescales differences in calcium gatting dynamics lead to the emergence of fundamentally new types of dynamics in the presence of in-vivo-like fluctuating input into a dendritic compartment.

**Fig. 1.**
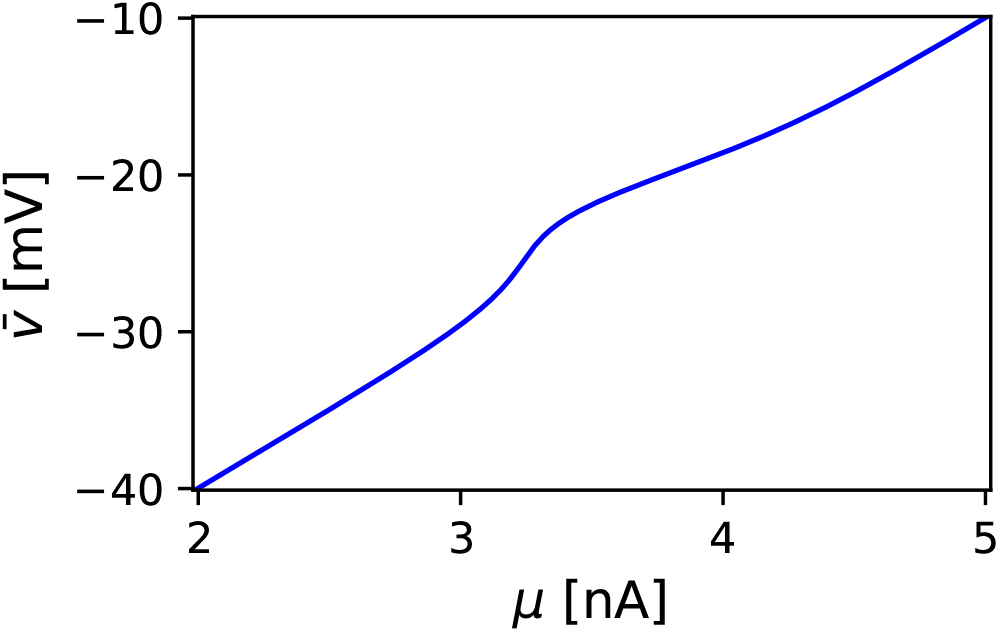
Input-output relation of a noiseless dendritic compartment. The equilibrium voltage (blue line), 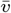, as a function of a constant input, *μ*, shows a monotonous relation.

## Noise-induced bistability with slow calcium activation and inactivation

The dendritic calcium dynamics can be mediated by gating variables whois their time-constants 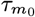 and 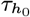 are much larger than the membrane time-constant *τ*_mem_. For these types of dynamics, in the timescale in which the voltage is fluctuating, we can assume both *m* and *h* are at their equilibrium 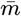 and 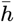, respectively. In this setting, the equilibrium dendritic membrane potential in Eq. (20), can be written as

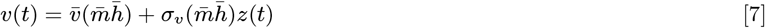

where 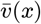 is given in Eq. (38),

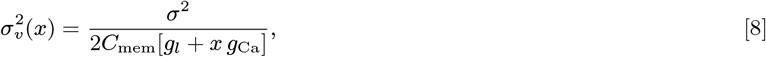

and *z*(*t*) is a Gaussian random variable with 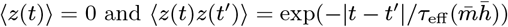, where

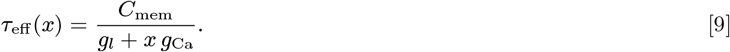

Since the calcium activation gating variable is slow 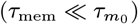, its *effective dynamic* is specified by

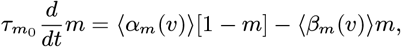

Thus, the equilibrium is given by

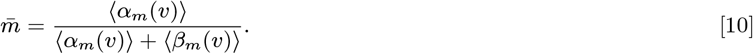

where

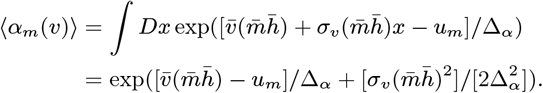

where 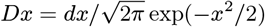 is the Gaussian measure. Similarly, 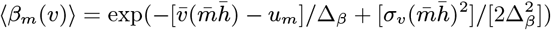. Analogously, it is straightforward to calculate the self-consistent average steady-state values for

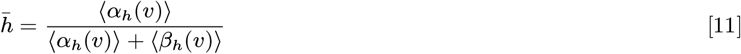

where, 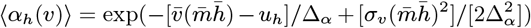 and 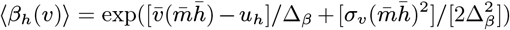. We plot these equilibrium values in Figure 2. In Figure 2.A, we demonstrate the input-output relation of the dendritic compartment for different values of noise. The result, following Figure 1, shows in the absence of noise (blue line in Figure 2.A), there is a monotonic increase in the average dendritic voltage 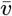 with small acceleration when calcium activation is sufficiently large. With small amounts of noise (green line in Figure 2.A), the acceleration increases, which corresponds to the shifts in the activation curve in Figure2.B and results in an increase in the maximum slope and level of 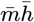(green solid and dashed lines in Figure 2.B). Notably, in the presence of sufficiently large noise (magenta line in Figure 2.A), we observe that 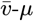 curve folds back, and there is a range of mean input for which multistability of 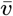 occurs. The critical value for noise level is given in the method section. In this region, three possible values for 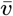 emerge for one input level. The middle point is an unstable fixed-point of the dynamics, and the two surrounding points are stable fixed-points for the equilibrium average dendritic voltage. With additional noise, this region of bistability shifts to the left, but its width saturates. To understand the primary mechanism for the emergence of bistability in this regime, in Figure 2.B, the values of 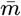 (dashed lines) and 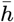 (solid lines) are plotted against 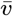, for different noise levels. In the absence of noise, the activation curve is sigmoid, where 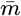 increases from zero to one as 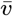 increases (blue dashed line in Figure 2.B). Similarly, the inactivation variable 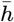 is a decreasing sigmoid (blue solid line in Figure 2.B). We also show half-potential values for the activation, *u_m_*, and inactivation, *u_h_*, by vertical gray dashed and solid lines. It is noteworthy that in the absence of noise, values of *u_m_* and *u_h_* are relatively close, and 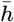 already starts to decrease when 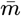 reaches a significant non-zero value. As a result, when *σ* = 0.0, the maximum value of 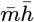 is also small. It also is important to note that the solution for 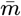 and 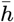 indicate that that adding noise to the input is equivalent to shifting *u_m_* to 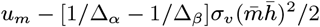 and 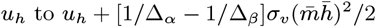. Therefore, if Δ*_α_* < Δ*_β_* then the value of 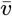, where the calcium channel activates (*m >* 0), shifts to left when the noise is increased, while the value of 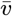, where the channel starts its inactivation (*h <* 1), shifts to the right. In the setting of Figure 2.B, with increasing noise, the activation equilibrium 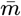 shifts to the left (green and magenta dashed lines in Figure 2.B) and inactivation equilibrium 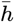 shifts to the right (green and magenta solid lines in Figure 2.B) and allows a higher amplitude in the *window current* of the gating variables into the dendrite. Accordingly, with sufficiently strong noise, the maximum value of 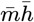 comes close to one and the maximum slope of 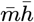 as the function of 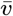 also significantly increases, which corresponds to the folding back phenomenon in the input-output relation in Figure 2.A. Therefore, sufficient noise determines the average dendritic voltage’s bistability as a function of mean input.

**Fig. 2.**
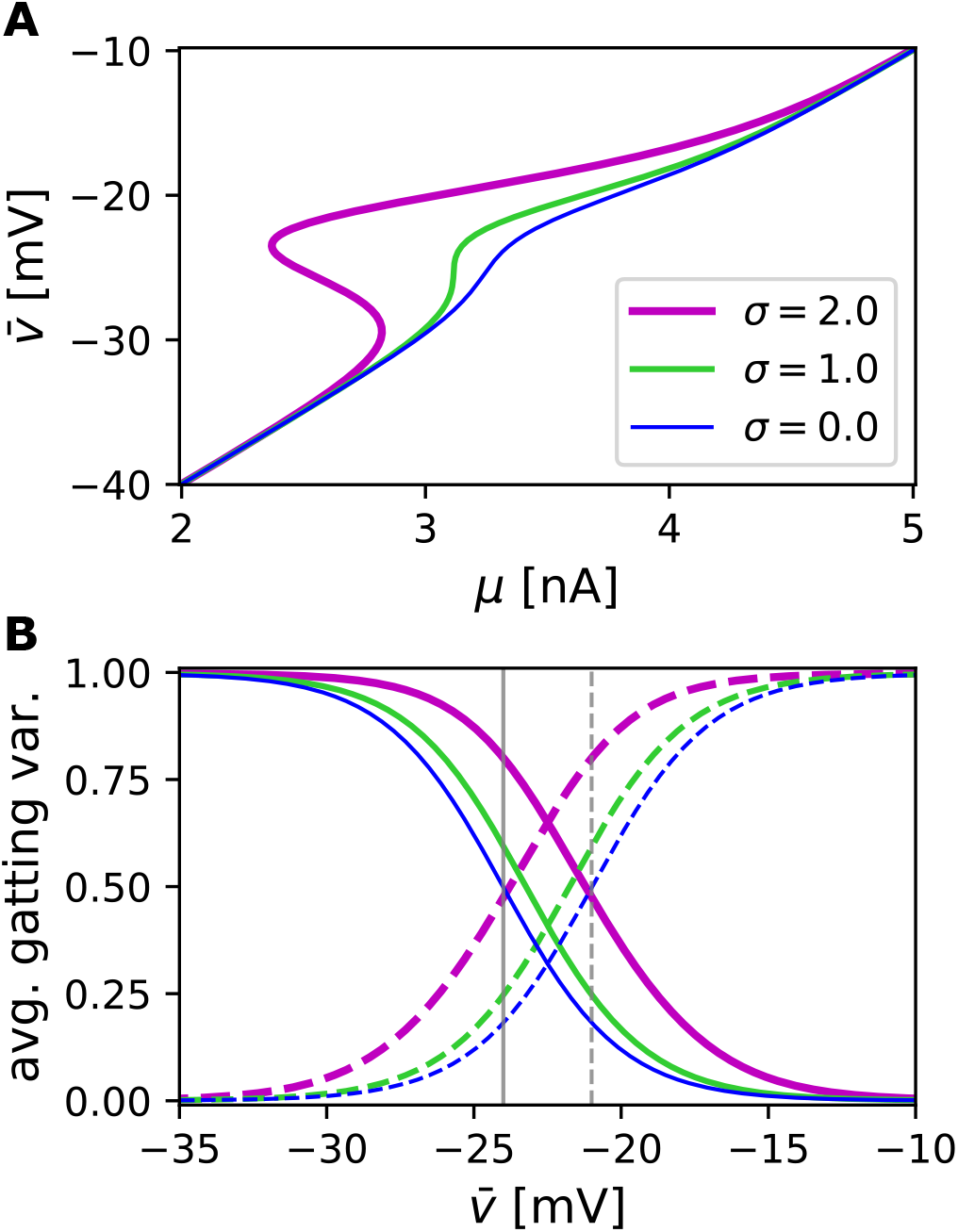
Noise-induced bistability in the dendritic input-output relation. (A) The equilibrium averaged voltage, 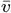, is plotted against mean input, μ, for different noise levels *σ* = 0.0 (blue line), *σ* = 1.0 (green line), and *σ* = 2.0 (magenta line) for a dendritic compartment with slow activation and inactivation of the calcium feedback. (B) The average equilibrium value of the calcium activation gating variable (dashed lines), 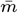, and inactivation gating variable (solid lines), 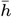, are plotted as functions of the equilibrium averaged voltage, 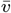, for different noise levels, corresponding to curves as color-coded in (A). The half-activation, u_m_, and -inactivation, uh, potentials in the deterministic system are indicated by vertical gray dashed- and solid-lines.

## Noise-induced non-monotonicity with fast calcium activation and slow inactivation

A major class of dendritic calcium channels exhibits a fast activation, while the inactivation variable is slower than membrane dynamics [13], 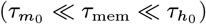, which allows the application of the slow-fast analysis. In the dendritic compartment with these type of calcium channels, we have the activation gating variable converges quickly to its equilibrium value at a given voltage, thus *m* is *m*_∞_(*v*) = *α_m_*(*v*)*/*[*α_m_*(*v*)+*β_m_*(*v*)], and in the timescale in which the voltage is fluctuating, *τ*_mem_, we can neglect the fluctuation in *h* around its equilibrium value 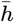, since 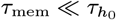. Hence, the dendritic compartment membrane potential dynamics in Eq. (20) becomes

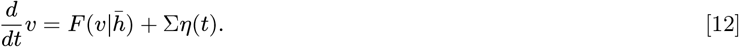

where 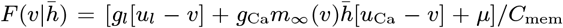 and Σ = *σ/C*_mem_. The Fokker-Planck equation [16], which describes the evolution of voltage distribution for Eq. (12), is given by

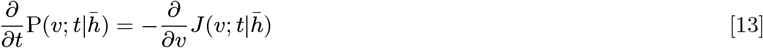

where 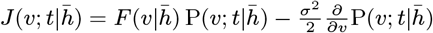. Thus, the equilibrium distribution of the voltage is

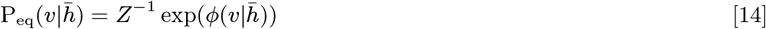

where 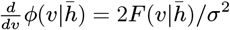 and *Z* is a constant that ensures 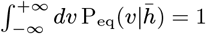. In this regime, the effective inactivation by

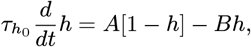

where 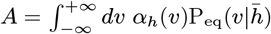, and 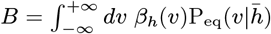. Thus, the self-consistent equilibrium value of 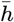 is specified by 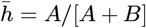, and the average equilibrium value of the activation gating variable is given by 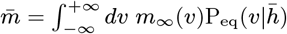. In Figure 3.A, we plot the equilibrium mean dendritic voltage 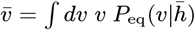 against *μ* for differe−n∞t values of *σ*. In the deterministic case, where *σ* = 0, we observe the average denetric voltage in response to an increase in *μ* is monotonously increasing, consistent with Figure 1. Interestingly, with sufficient noise, non-monotonous behavior of 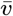 emerges. In a certain range, we observe that an increase in *μ* reduces the corresponding 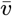 (green and magenta lines in Figure 3.A). Thus, in the presence of in-vivo-like fluctuations, we observe a noise-induced non-monotonicity in the canonical dendritic compartment model with fast activation and slow inactivation of the calcium gating variables. To illustrate the underlying mechanism for this noise-induced nonlinearity in this regime, we also plot both equilibrium averages of 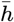 and 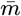 as the function of 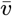 for different levels of noise in Figure 3.B. Figure 3.B shows that the activation and inactivation are standard sigmoidal curves in the absence of noise (blue dashed- and solid-lines in Figure 3.B). In Figure 3.B, as a reference point, we indicate half-activation potential, *u_m_*, and half-inactivation potential, *u_h_*, by the vertical gray dashed- and solid-lines, respectively. By increasing the noise level, 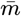 starts to increase in the lower voltages and is tightly coupled with the dendritic average voltage (green and magenta dashed lines in Figure 3.B). An increase in the noise level shifts the inactivation equilibrium to the left (magenta solid-line in Figure 3.B) and when 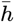 starts to decrease, it causes a reduction on the average voltage as well (green and magenta solid lines in Figure 3.B). In return, the positive voltage feedback modifies the slope of the averaged inactivation in such a way that we can find three equilibria values for 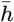 (green solid-line in Figure 3.B). This determines also a strong nonlinearity on 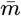 during the reduction of 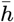 (green and magenta solid lines in Figure 3.B), which causes the corresponding non-monotonous behavior in 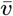. Hence, these noise-induced nonlinearities contribute to the corresponding non-monotonous behavior in the input-output relation of the dendritic compartment in Figure.3.A.

**Fig. 3.**
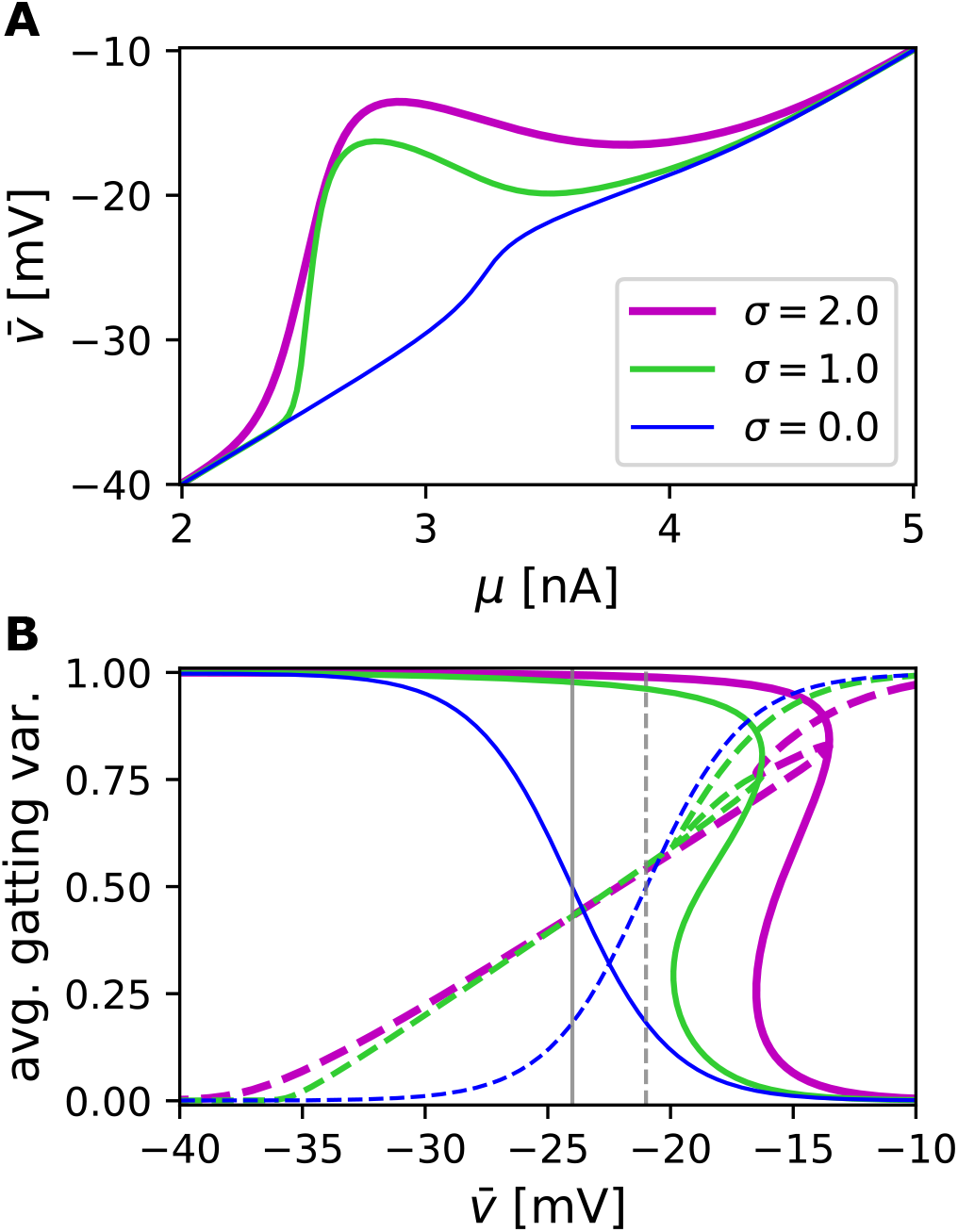
Noise-induced non-monotonicity in the dendritic input-output relation. **(A)** The equilibrium averaged voltage, 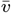, is plotted against mean input, *μ*, for different noise levels *σ* = 0.0 (blue line), *σ* = 1.0 (green line), and *σ* = 2.0 (magenta line) for a dendritic compartment with a fast activation and slow inactivation of the calcium feedback. **(B)** The average equilibrium value of the calcium activation gating variable (dashed line), 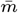, and inactivation gating variable (solid lines), 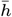, are plotted as functions of the equilibrium averaged voltage, 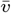, for different noise levels, corresponding to curves as color-coded in (A). The half-activation, *u_m_*, and -inactivation, *u_h_*, potentials in the deterministic system are indicated by vertical gray dashed- and solid-lines.

## Noise-induced switching in the bistable regime

The emergence of noise-induced phenomena in Figures 2 and 3 indicate that in-vivo-like noise significantly modifies the dendritic input-output relation. The non-monotonicity in Figure 3 suppresses input fluctuations. Therefore, correlations in the voltage decay on the timescales at the closest to *τ*_mem_. However, noise-induced bistability can enhance the system’s sensitivity to small changes, and it can modify the timescales of the system. In deterministic bistable systems, the additive noise allows switching between the possible solutions with a particular time-constant [17]. However, noise induces the bistability here, and contrary to the assumptions in the slow-fast analysis (i.e., 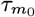 and 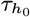 are infinitely large), the realistic values of 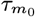 and 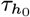 modifies the dynamics. To investigate the effect of finite 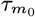 and 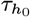 we simulate the full system given in Eq. (20), Eq. (23) and Eq. (24). In Figure 4.A, we show a response of membrane voltage *v* at its equilibrium to a short perturbation in the mean input *μ*, when *σ* = 0.0 (blue line in Figure 4.A). The brief positive perturbation activates calcium followed by inactivation. The induced dynamics capture a phenomenon that is widely known as a dendritic calcium spike [5]. The voltage trace of the deterministic system (blue line in Figure 4.A) goes back to the original equilibrium value with the timescales of 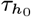. We further show the voltage response averaged over 5 104 realizations of the noise, for small and large *σ*. In the case of small noise (green line in Figure 4.A, *σ* = 0.4), we observe the response to the perturbation goes back to the deterministic baseline with the same timescale as the noiseless system. This result is consistent with the lack of multistability when *σ* = 0.0 and *σ* = 0.4 (blue and green lines in Figure 4.A). In contrast, with sufficient noise level that induces the bistability, the response to the perturbation decays much slower than the deterministic case (blue line in Figure 4.A) and the system with small noise (green line in Figure 4.A). It is apparent that when 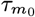 and 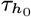 are finite, the voltage fluctuation also induces fluctuations in *m* and *h*. These fluctuations can occasionally cause switching between the two stable states predicated by the slow-fast analysis (Figure 2). To demonstrate this, we plot the average voltage (across trails) on a longer time window in Figure 4.B. We observe that on the timescales slower than 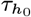, the average membrane voltage converges to an intermittent value between two predicted stable fixed points by the slow-fast analysis. The convergence to this intermittent value in the long timescale is independent of the perturbation as the system with the same level of noise without any perturbation also converges to this intermediate level of average voltage (black line in Figure 4.B).

**Fig. 4.**
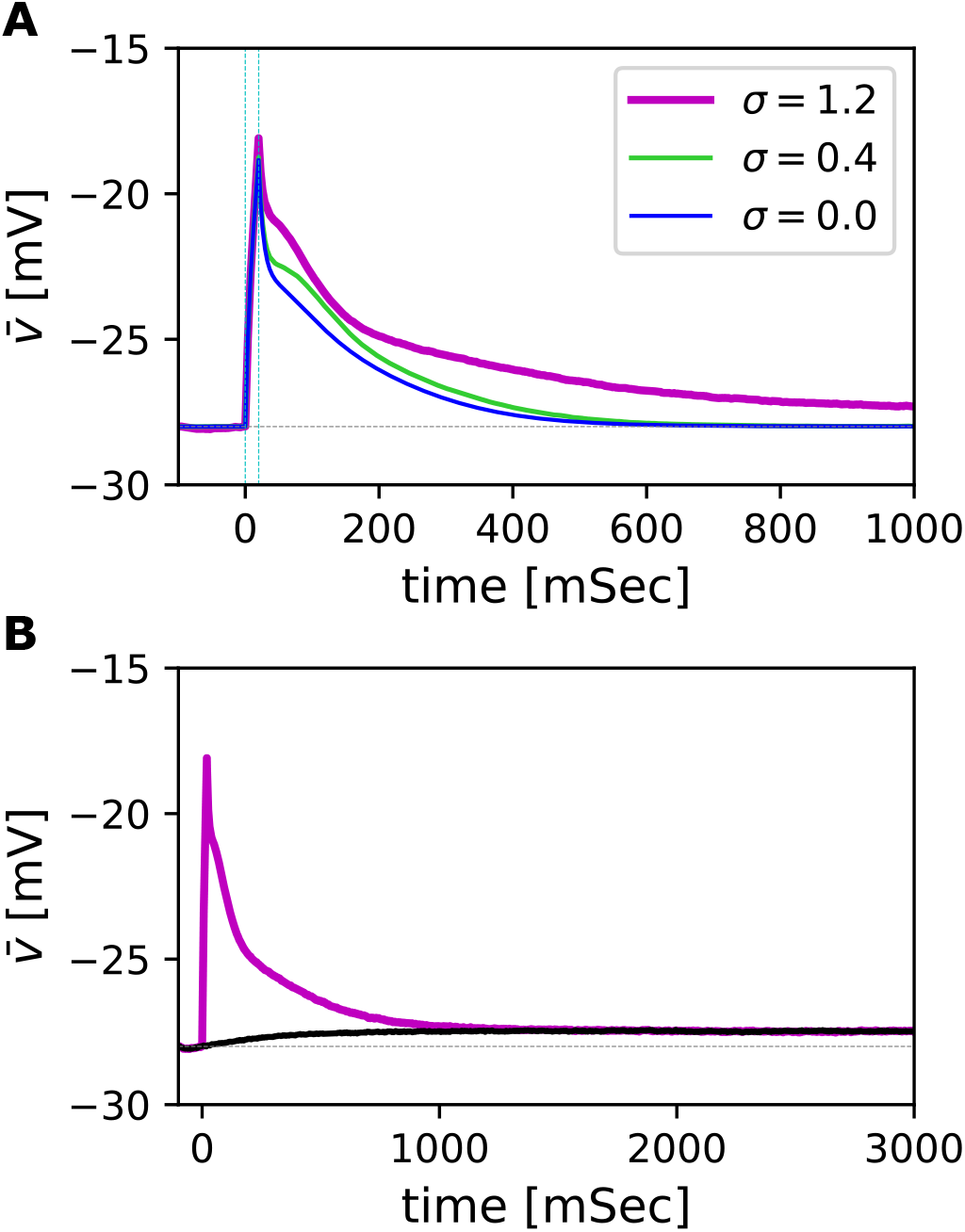
Simulation of a bistable dendritic compartment with a slow activation and inactivation of calcium feedback. **(A)** The dendritic voltage response of the full system to a short-lived perturbation (20 millisecond as it is indicated by vertical cyan dashed lines) in the mean input for different noise levels *σ* = 0.0 (blue line), *σ* = 0.4 (green line), and *σ* = 1.2 (magenta line), where 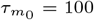 and 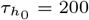. The voltage traces for the noise level *σ* = 0.4 (green line) and *σ* = 1.2 (magenta line) are averaged over 5×10^4^ trails. The baseline for the deterministic system (blue line) is indicated by a horizontal gray dashed line. **(B)**. The trial averaged voltage trace of 5×10^4^ simulation runs of the full system with a brief perturbation (magenta line) as in (A), and without the perturbation (black line) for the noise level *σ* = 1.2 that induces bistability. The system deviates from the initial condition at the lower stable solution (indicated by the horizontal gray dashed line) and settles slowly into an intermediate voltage value due to stochastic switches between solutions, irrespective of the perturbation.

The results in Figure 4 indicate that a complete description of the bistable stochastic dendritic dynamics may require the inclusion of the fluctuations in *v*, *m*, and *h* to understand the noise-induced phenomena and its timescales. This involves the translation of the state variables’ stochastic differential equations to a probability density evolution, using the Fokker-Planck approach [16]. This approach specifies an evolution equation for the state variables’ joint probability density, *v*, *m* and *h*, as P(*v, m, h*; *t*). However, the inclusion of all state variables does not lead to readily interpretable results. It does not use the separation of timescales between *τ*_mem_ and both 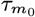, 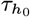, which allows one to determine the statistics of *v* and derive self-consistent stochastic differential equations for *m* and *h* on a timescale larger than *τ*_mem_ based on the voltage statistics. Additionally, it is also important to note that our results demonstrated in Figure 2 indicate that the emergence of bistability is initiated by the activation of the calcium channels. Therefore, the separation of timescales and the contribution of positive activation feedback suggest that a parsimonious approximate description of the dynamics can be developed. We take into account the fluctuations in *m* but neglect the fluctuation in *h*. This is equivalent to assuming 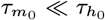. To this end, we assume 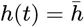 and it is in the equilibrium, and *m* does not change on the timescale of *τ*_mem_, therefore

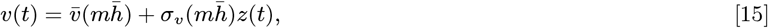

where *z*(*t*) is a Gaussian random variable with 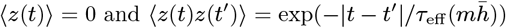 and 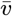, *σ_v_* and *τ_eff_* are given in Eq. (38), Eq. (39) and Eq. (40), respectively. In this setting, the fluctuation with strength 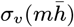 in the dendritic voltage induces fluctuation in *m*. On the 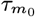 timescale, these fluctuations can be approximated by a white noise process, and therefore we obtain

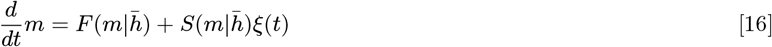

where *ξ*(*t*) is a standard white noise and 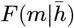 and 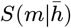 are chosen such that over a time-interval of *δ*, where 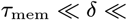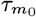, the Eq. (16) becomes approximately same as Eq. (23). In the method section, we derive 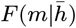 and 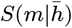 functions. The stochastic differential equation in Eq. (16) is also called the Langevin equation, 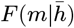 and 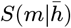 are drift and diffusion terms of the stochastic process. From this Langevin equation, we can readily derive a Fokker-Planck equation for the evolution of the probability density 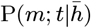 which satisfies

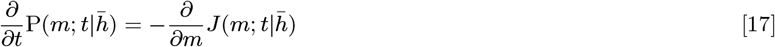

where 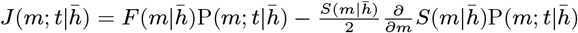 with the boundary condition of 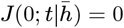. The stationary distribution, 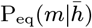, from the Fokker–Planck equation in Eq. (17) can be explicitly derived

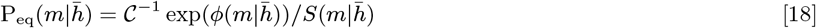

where 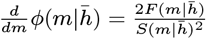 and 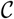 is a constant that ensures 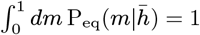. Now, we obtain the self-consistent solution,

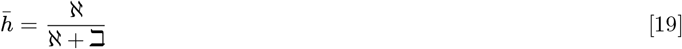

where 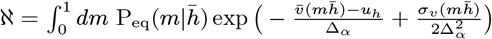 and 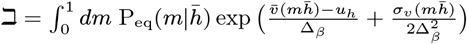. In Figure 5, we illustrate the equilibrium distribution as expressed in Eq. (18) for three values of mean input and *σ* = 2.0. We show for a value of *μ* below the bistable region, the probability mass is concentrated close to zero (blue dashed-dotted line in Figure 5). It is also shown that for a value of *μ* beyond the bistable region, the density peaks significantly close to one (magenta dashed line in Figure 5). However, for an intermittent value of *μ*, the probability density becomes pronouncedly bimodal, reflecting bistable behavior in the stochastic dynamics of *m* (black solid line in Figure 5). In this case, noise disperses the level of *m* around the two peaks, corresponding to the bistable solutions. Therefore, by initializing the system near one of the peaks, the stochastic dynamics result in a relatively rapid fluctuation around that peak value. On a longer timescale, the noise promotes flights to the other peak value and therefore induces stochastic switching between peaks. The timescales on which this happens can be inferred from the eigenvalue spectrum of the Fokker-Planck operator [16]. We numerically calculate the Fokker-Planck operator’s eigenvalue spectrum corresponding to Eq. (17) (see the methods for details). The largest eigenvalue of the Fokker-Planck operator *λ*0 = 0, whose eigenfunction corresponds to the equilibrium distribution 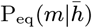; since this solution is stable, all other eigenvalues have negative real parts. The rate that the system approaches this equilibrium distribution from an arbitrary initial condition is determined by the second eigenvalue, *λ*1, of the Fokker-Planck operator. The timescale over which the probability density of *m* approaches equilibrium is given by the relaxation time, 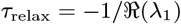. In Figure 6.A, we illustrate *τ*_relax_ as a function of *σ*, for a value of *μ* corresponding to 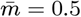 in Eq. (41). Figure 6.A shows the relaxation time increases as sufficient noise induces bistability and reaches its maximum, thereafter it decays gradually to *τ*_mem_ as additional noise facilitates faster switches between two solutions. Furthermore, in Figure 6.B, we plot *τ*_relax_ against *μ* for *σ* = 1.5 ensuring that the system exhibits bistability (black line in Figure 6.B). The relaxation time (black line in Figure 6.B) reaches its maximum for *μ* in the bistable regain. Note that the relaxation time (black line in Figure 6.B) substantially deviates from the deterministic system (blue line in Figure 6.B). In Figure 6.B, we also calculate the full system autocorrelation timescales of *m* fluctuations, by simulating dynamics in Eq. (20), Eq. (23) and Eq. (24) for two different values of 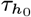 (black and green cross symbols in Figure 6.B). The relaxation time derived from the parsimonious approximate description of the system in Eq. (17) (black line in Figure 6.B) captures the behavior of the full system dynamics when *τ*_relax_ 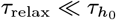 (black cross symbols in Figure 6.B, where 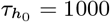). However, it becomes apparent that *τ*_relax_ derived from the one-dimensional Fokker-Planck approximation in Eq. (17) can be of the same order of 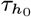 (green cross symbols in Figure 6.B, where 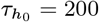). Therefore, the one-dimensional approximation in Eq. (17) (black line in Figure .B) cannot quantitatively predict the timescales of the full system’s fluctuation. In this case, we require the inclusion of fluctuations in *h*, by constructing a two-dimensional Fokker-Planck operator for the evolution of the joint probability density *mathrmP* (*m, h*; *t*), using the same approach as Eq. (17) (see the the methods). In this case, the relaxation time calculated from the approximate two-dimensional Fokker-Planck operator description of the system P(*m, h*; *t*) (green line in Figure 6.B) becomes close to the estimated autocorrelation timescales of the full system (green cross symbols in Figure 6.B)

**Fig. 5.**
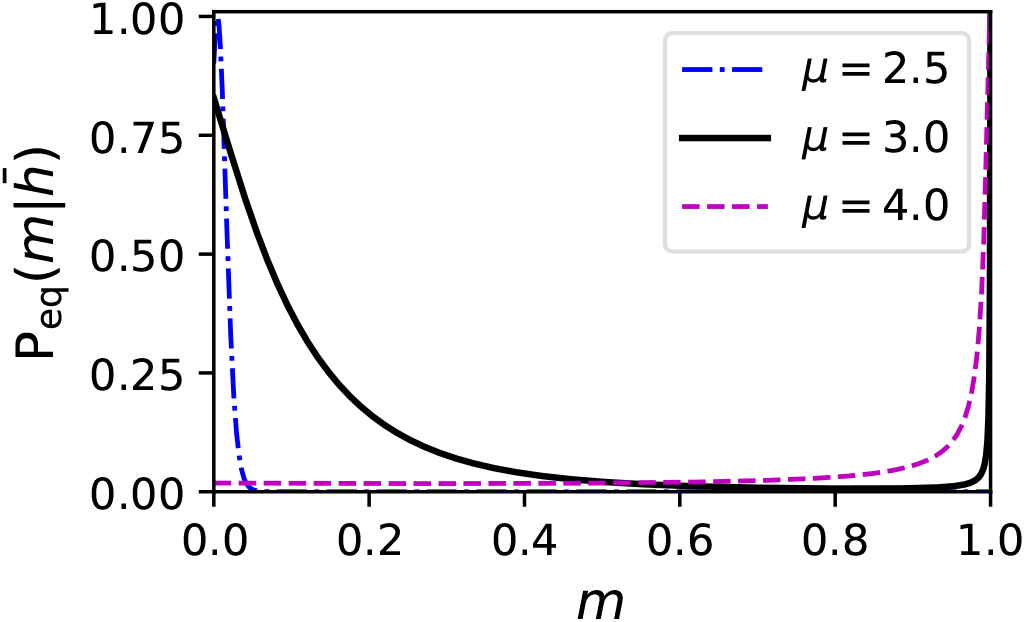
Equilibrium distribution of the activation gating variable in the bistable dendritic compartment. The equilibrium density of *m* is given by Eq. (18) for different values of mean input below the bistable region (blue dashed-dotted line; *μ* = 2.5, in the bistable regain (black solid line; *μ* = 3.0), and beyond the bistable regain (magenta dashed line; *μ* = 4.0), for *σ* = 2.0 and 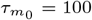. The maximum values are normalized to one.

**Fig. 6.**
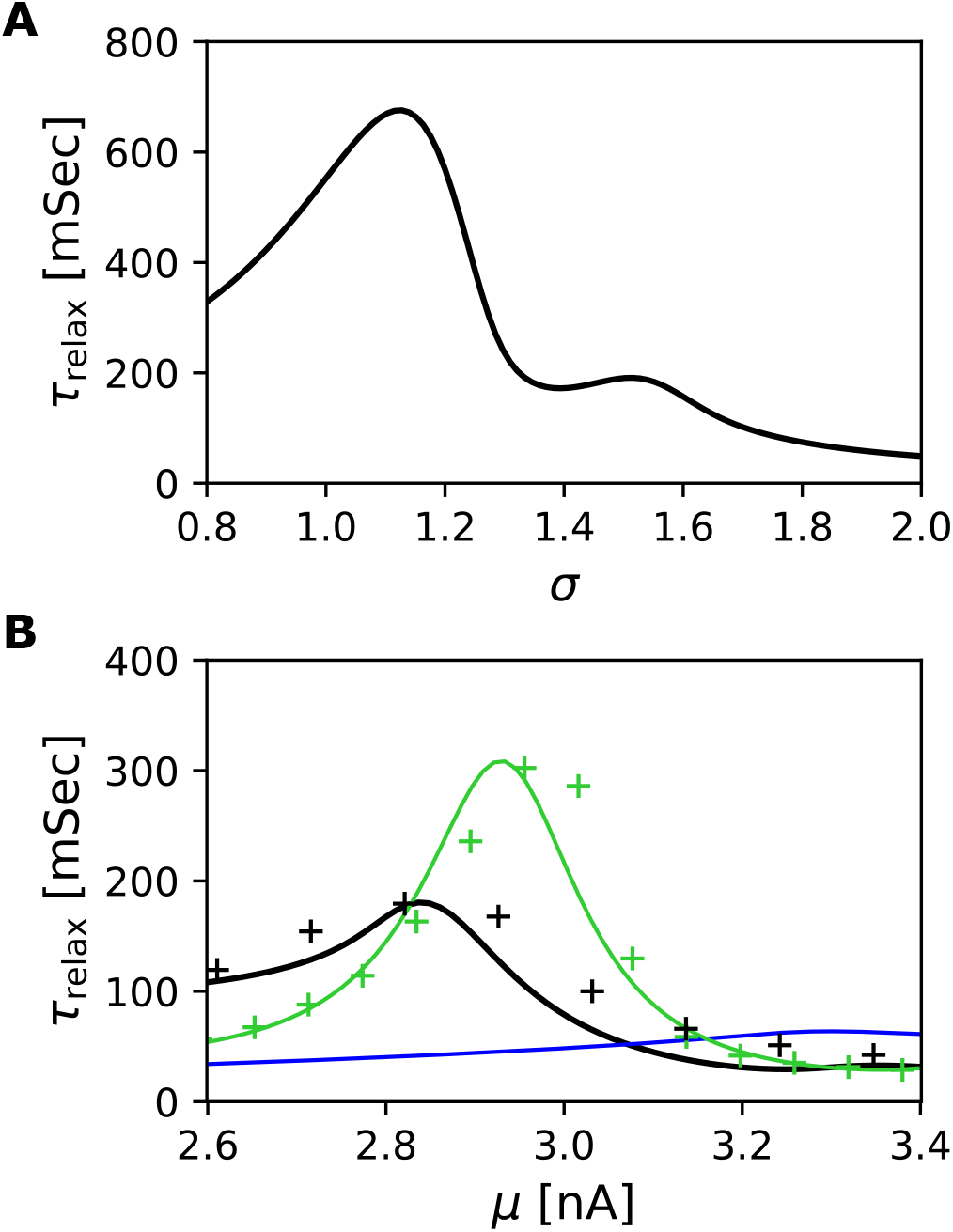
The timescale of the fluctuations in the bistable dendritic compartment. **(A)** The relaxation time, *τ*_relax_, is plotted as a function of the noise level, *σ*. We adjust mean input, *μ*, for various noise levels to ensure that 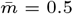, and therefore the system stays in the bistable region. We choose 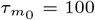. **(B)**. The black line shows the relaxation time derived from the one-dimensional Fokker-Planck operator for *m* is plotted against mean input for noise level *σ* = 1.5 and 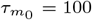. Corresponding to the black line, the autocorrelation timescales of *m* fluctuations estimated from the simulations, when 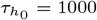, are shown with black crosses. The green line represents the relaxation time derived from the two-dimensional Fokker-Planck operator for the evolution of joint distribution of *m* and *h* (see the methods), as a function of mean input, where *σ* = 1.5, 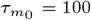 and 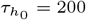. Corresponding to the green line, the autocorrelation timescale of *m* is estimated from simulations are shown with green crosses. The blue line is the effective timescale of *m* in the deterministic system.

## Discussion

The diverse calcium machinery of dendrites has been studied intensively in the last three decades [1]. While dendritic calcium channels exhibit enormous heterogeneity, their dynamics are generally known to be slow [13, 14]. In the absence of noise with constant input, the differences in the dynamics of calcium gating variables do not contribute to the dendritic input-output relation (Figure 1). Our analytical delve into an active dendritic compartment uncovers that the differences in the dynamics of calcium gatting variables leads to qualitatively different types of noise-induced phenomena in in-vivo-like conditions. Our results show that in the case of a fast activation of the calcium gating variable, noise-induced non-monotonicity arises (Figure 3). For the case where both activation and inactivation gating variables are slow, noise-induced bistability emerges in the dendritic compartment′s input-output relation (Figures 2 and 5). The noise-induced non-monotonicity facilitates the shaping of input fluctuations by suppression of fast changes, and an interesting byproduct of noise-induced bistability is the introduction of a long timescale due to stochastic switches between two solutions. The switching timescales can be calculated, using the approximate Fokker-Planck equation (Figure 6).

These noise-induced phenomena can have functional importance in the single neuron strategies to integrate fluctuating input by its various active dendritic channels. Our results show that dendrites could adjust their input-output relation in response to noisy input to maintain different sensitivities to changes. It is important to note that here we study the dynamics of an isolated dendritic compartment with an in-vivo-like fluctuation in the input. A necessary future research step is to investigate interactions among dendritic compartments and their coupling with soma to understand how fluctuating dendritic inputs affect the neuronal activity and network dynamics. For example, pyramidal neurons in cortical layer 2-3 networks receive their feed-forward input via their basal dendrites, and the apical dendrites collect recurrent projections [18, 19]. Therefore, a differential projection of somatostatin (SOM)-positive and parvalbumin (PV)-positive interneurons recurrent input into the apical dendrites and various noise-induced nonlinearities could be used to modify the effect of feed-forward and recurrent input of the network dynamics [20, 21].

We conclude that noise can function as an organizing principle that leads to new kinds of dynamics in the neuronal dendritic compartment. Our results add to previous studies that noise induces order in many other biological and physical systems [8–11] and imply that noise-induced dynamical processes can be an underlying mechanism for robustness in systems operating in a fluctuating environment.

## Methods

We study the influence of noisy external input in Eq. (20) on the nonlinear feedback of voltage-dependent calcium channels given in Eq. (23) and Eq. (24). We first perform the slow-fast analysis exploiting the timescales differences in the system as described in the main text. Below, we further calculate critical values for *g*_Ca_ and *σ* in the case of the emergence of noise-induced bistability. Next, we describe the stochastic switching phenomenon due to finite timescales of calcium dynamics in the bistable system (Figure 4). To calculate the switching time between the stable solutions, we use the system’s Fokker-Planck operator eigenvalue decomposition for the activation variable, *m*, as it is described in the main text. The complete derivation of Fokker-Planck approximation in Eq. (17) and the numerical method to calculate its corresponding switching time, *τ*_relax_, are provided below. To achieve a better quantitative approximation for the cases that timescales of activation and inactivation are close, we use a two-dimensional Fokker-Planck equation for the joint evolution of *m* and *h* (Figure 6.B). The underlining two-dimensional Fokker-Planck description in Figure 6.B is given in the following section.

### 1. The emergence of meta-stability in the slow-fast analysis

To recapitulate, in this paper we consider the canonical conductance-based model of a dendritic compartment as expressed by

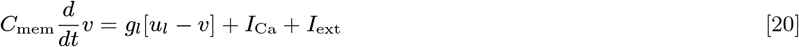

where, *C*_mem_ and *g_l_* are the membrane capacitance and leak conductance per unit area, *u_l_* is the leak reverse potential, and *I*_Ca_ is the active dendritic feedback. We model in-vivo-like external input per unit area, *I*ext, by a generic white-noise process as

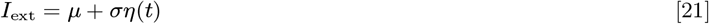

where, *μ* and *σ* are the mean and standard deviation of the process, and *η*(*t*) is a Gaussian white noise variable, where 〈*η*(*t*)〉 = 0 and 〈*η*(*t*)*η*(*t′*)〉 = δ(*t* − *t′*). We use the notation of 〈.〉 to denote averaging over the noise realization. The dendritic active feedback properties due to calcium dynamics in Eq. (20) is captured by

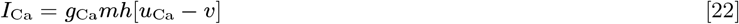

where, *g*_Ca_ and *u*_Ca_ are the maximum calcium conductance per unit area and its reverse potential, respectively. The dynamics of the activation gating variable, *m*, and inactivation gating variable, *h*, are given by

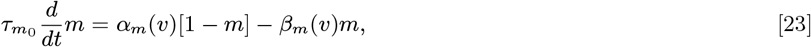

and

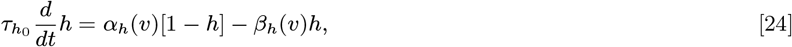

respectively. First, we look at the steady-state solution, 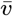, in the noise-less system (*σ* = 0), which it satisfies

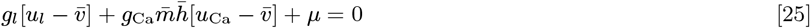

where, 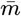 and 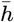 are the steady-state for activation and inactivation, respectively, and are given by

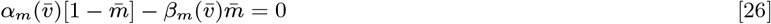

and

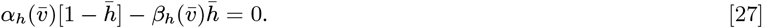

Thus, we haves

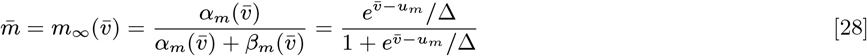

and

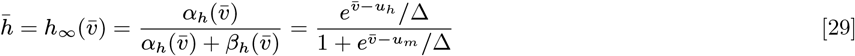

where 1/Δ = 1/Δ*_α_* + 1/Δ*_β_*. Therefore, we can write

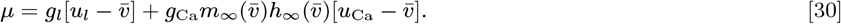

This indicates 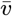 varies *monotonously*. Next, we investigate under which condition the solution is mono-stable. To do so, we parametrize 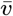 and *μ* as

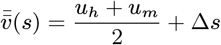

and

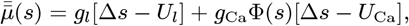

where *U_l_* = *u_l_* − [*u_m_* + *u_h_*]/2, *U*_Ca_ = *u*_Ca_ − [*u_m_* + *u_h_*]/2, and

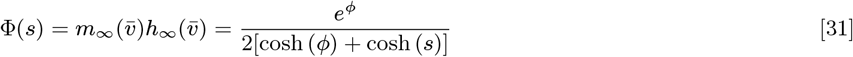

where 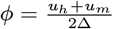. Since 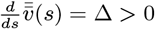, the system is mono-stable if and only if

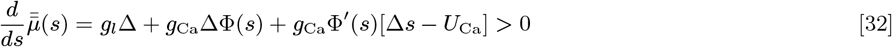

for all *s*. To verify this condition, we require to find a minimum point and investigate its sign.The system has its minimum at *s* = *s*_min_, which satisfies

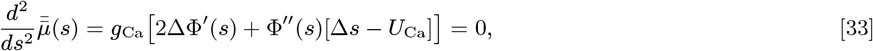

and hence *s*_min_ is given by

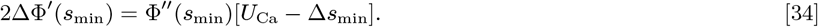

Hence, the input-output relation is *mono-stable* if and only if

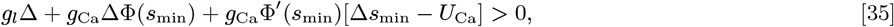

which equivalently implies

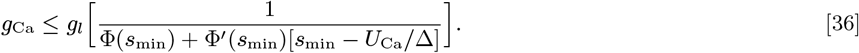

This condition determines a critical value of calcium conductance, 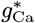, where below which input-output relation is *mono-stable* and above which it is *multi-stable*. Our analysis shows that the system is mono-stable when *u_h_ u_m_* is comparable to Δ or smaller (keeping the other parameters fixed).

Now, we show the introduction of noise (*σ >* 0) effectively lower the critical value of calcium conductance, 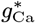, in a regime that both activation and inactivation dynamics are much slower than membrane potential fluctuation. In the noisy system, where 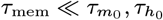, we can assume 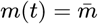 and 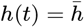. Therefore, the average equilibrium voltage satisfies

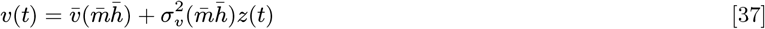

where 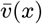 and *σ_v_* (*x*) are defined as

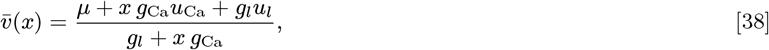

and

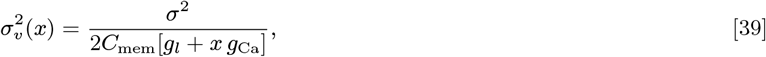

and *z*(*t*) is a Gaussian random variable with zero mean and temporal correlation given by 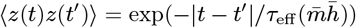,

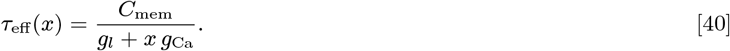

The self-consistency allows us to determined 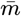 as

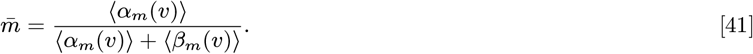

where

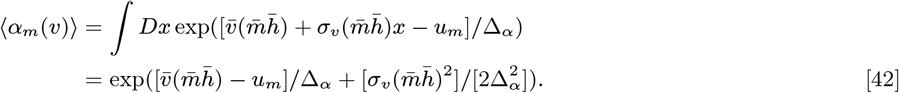

where 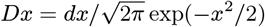 is the Gaussian measure. Similarly, 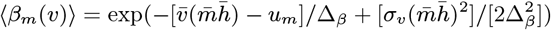. Analogously, it is straightforward to calculate the self-consistent average steady-state values for

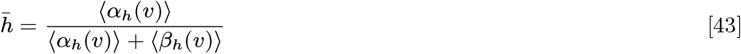

where, 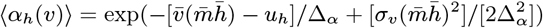 and 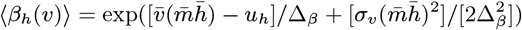. Now, we assume Δ*_β_* = *κ*Δ*_α_* (or Δ*_α_* = Δ/[1 + *κ*]) and Δ*_β_* = Δ[1 + *κ*]. Thus, we obtain

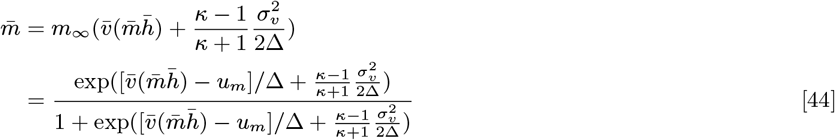

and analogously 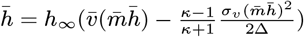. Using parameterization

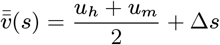

and we have

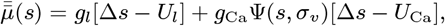

where,

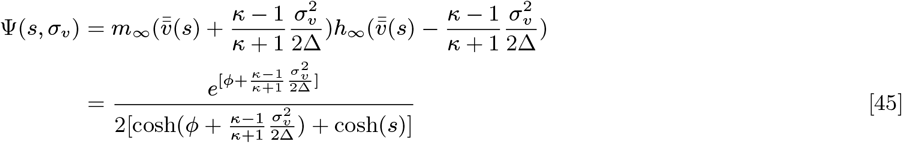

where 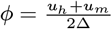 as in Eq. (31). Therefore, adding noise has the effect of shifting *φ* to 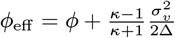 and for *κ>* 1 input noise increases *phi*_eff_. As a result, the critical value of calcium conductance, 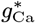 is a decreasing function of *φ*_eff_, denoted as 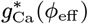, and as *φ*_eff_⟶∞, the value of 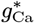 converges to its asymptotic minimum 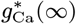. Thus, if *g*_Ca_ is such that 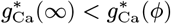 the dendrite input-output relation is mono-sable in the absence of the noise. However, there is a critical noise level, *σ*\*, such that at 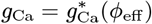 the system switches form a mono-stable to a *bistable* input-output relation.

### 2. The approximate Fokker-Planck description of the system

#### One-dimensional Fokker-Planck equation for the evolution of slow activation

The slow-fast analysis for the dendrite compartment predicts that when 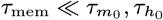, the noise-induced bistability in the input-output relation of the dendrite. However, in the case of finite but large time-constants for calcium gating variables, the noise also facilitates switching between solutions. To understand this switching phenomenon, we study the approximate solution of the probability evolution of the system, in the case where 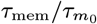 and 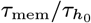 are small and finite. First, we will investigate the case where 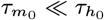 and thus we can neglect the fluctuations in *h*(*t*) and assume that 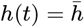. Moreover, since 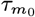 is much larger than *τ*_mem_, the statistics of the voltage in the equilibrium to the leading order is given by,

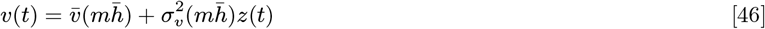

where *z*(*t*) is a centred Gaussian variable with 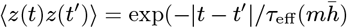. To describe the dynamics of *m*, it is useful to rescale time 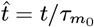, thus

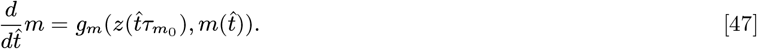

where 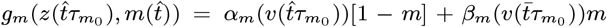, where *v* as it is defined in Eq. (46). Assuming that *m*(0) = *m*0, we want to derive the contribution of rapid fluctuations in *v* to the distribution of *m*, in a time window 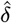, where 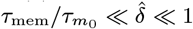. Hereafter, to simplify notation we define 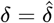. The average displacement can be calculated by

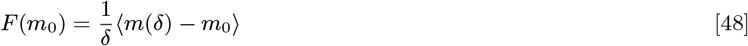

and its corresponding variance specifies by

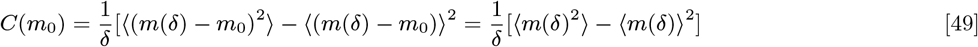

We have

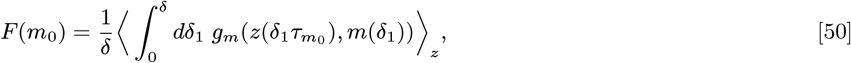

where 〈〉_*z*_ is the averaging operator over the Gaussian variable *z*. Since *m*(*δ*) close to *m*_0_, we Taylor expand the right-hand-side (RHS) of the above equation around *m*_0_ and write

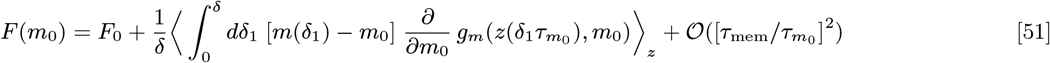

where 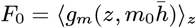 and it corresponds to the solution of 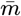 in the slow-fast analysis. The second term is an additional contribution to the drift results from fluctuations around that solution due to noise in the voltage value. We show below that this additional contribution to the drift is of 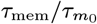 and the higher order terms are at most of the order of 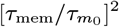. It should be also noted that *m*(*δ*_1_) and 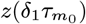, inside the integral (the second term), cannot be assume to be independent and therefore to the leading order, furthermore

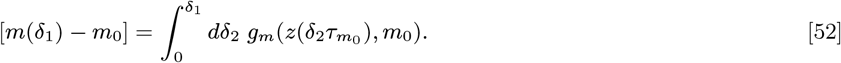

Thus, we obtain

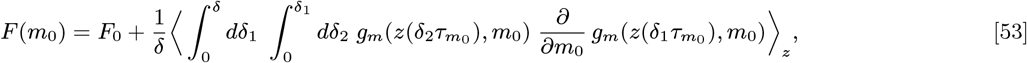

Note that the joint distribution of noise variables *z*(*t*_1_) and *z*(*t*_2_), P(*z*(*t*_1_), *z*(*t*_2_)), only depends |*t*_1_ − *t*_2_|, therefore in general for two arbitrary function of ℸ and ℶ we have

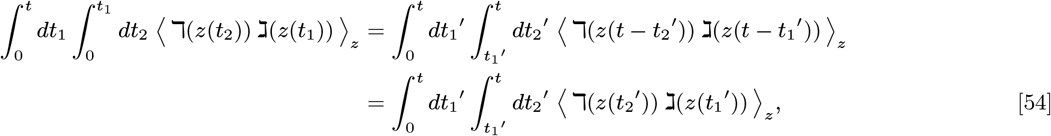

using change of the variables *t*_1_’ = *t* − *t*_1_ and *t*_2_’ = *t* − *t*_2_. Thus, this time-invariant properties implies

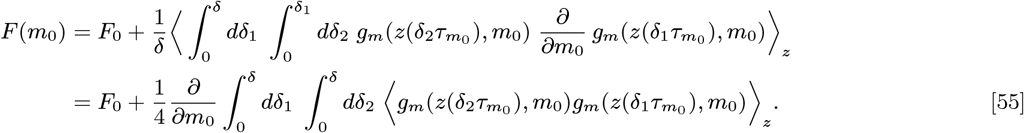

The integral in the above equation interestingly corresponds to the diffusion term, *C*(*m*_0_). Following the definition in Eq. (49), to the leading order expansion

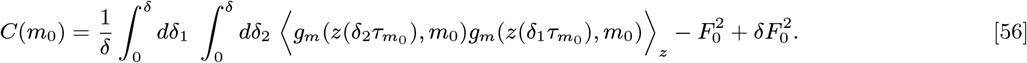

By using *δ*_−_ = *δ*_1_ − *δ*_2_, *δ*_+_ = *δ*_1_ + *δ*_2_, and *dδ*_−_*dδ*_+_ = 2*dδ*_1_*dδ*_2_, thus

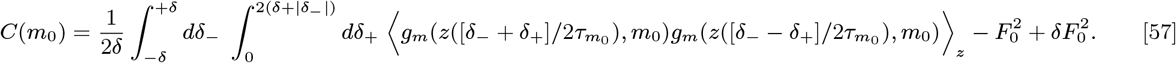

Because

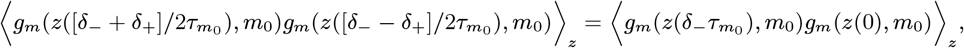

and the symmetry in the noise variable auto-correlation in Eq. (54), we can write

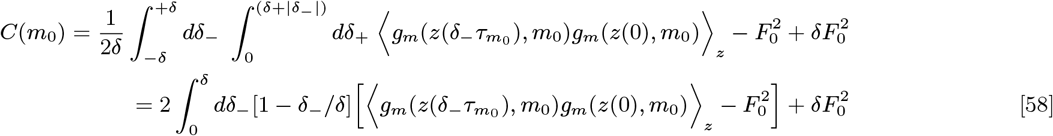

Note that the term in the bracket in the integrand becomes negligible for 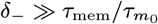 and thus we can neglect *δ*_−_*/δ* in the regime of slow dynamics for *m*. The integrand is also negligible for for large *δ*_−_, hence we can change the upper bounds of the integration

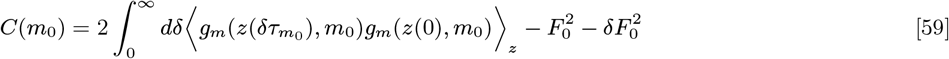

Here, we take *δ* → 0 thus

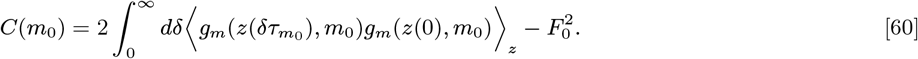

Therefore, the corresponding Fokker-Planck equation of the system with the drift as in Eq. (55) and the diffusion as in Eq. (60) is

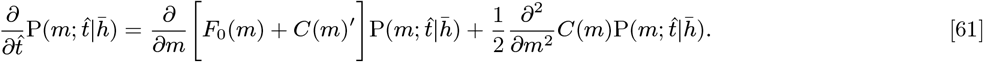

This is the Fokker-Planck equation that naturally emerges as it is expected in the Stratonovich approach. Note that in the main text, the time-rescaled Langevin equation is given by

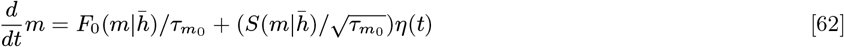

where 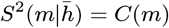,

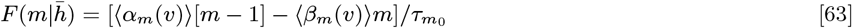

and

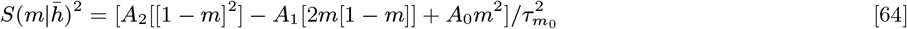

where,

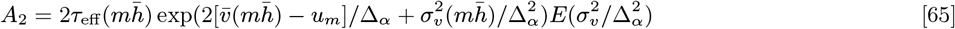

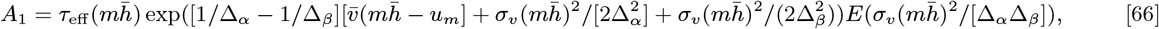

and

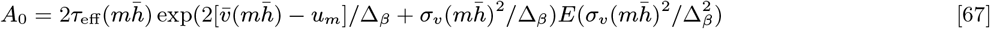

Here, we have 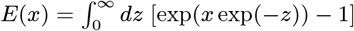. Note that in Eq. (38), we assume the system is in statistical equilibrium. Naively, this assumption seems incorrect, however, the higher order corrections of 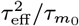 and therefore do not contribute to the calculation presented here.

#### Two-dimensional Fokker-Planck equation for evolution of slow *m* and *h*

In the condition where the timescale of inactivation, 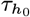, is comparable to 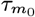 or the relaxation time of the system, we need to include the fluctuations of both *h* and *m* in the description of stochastic dynamics in the form of two-dimensional Fokker-Planck equation. To calculate the Fokker-Planck equation for the joint distribution of *m* and *h* we use the same approach as above. We re-scale the dynamics of *m* and *h* as

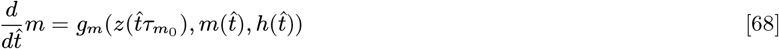

and

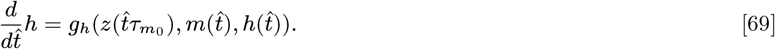

and therefore the corresponding drifts defined as

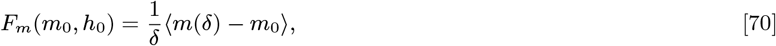

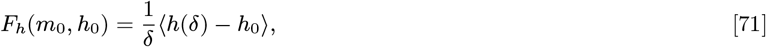

and the corresponding diffusion factors specifies by

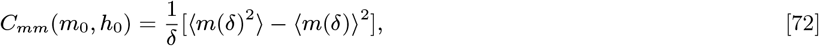

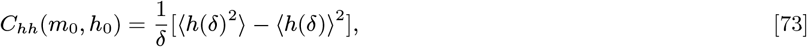

The co-variation between *m* and *h* is defined as

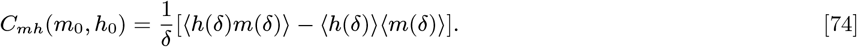

The derivation of two-dimensional Fokker-Planck equation follows as above, which it can be written as

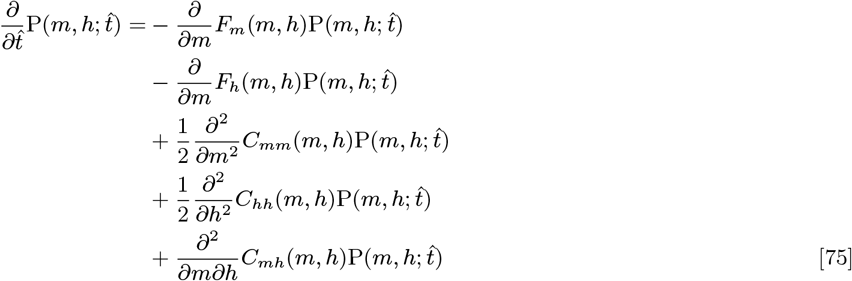

To keep the notation compacted, we define

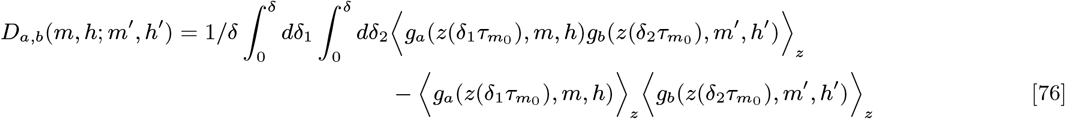

where *a* ∈ {*m, h*} and *b* ∈ {*m, h*}. Therefore, *C_mm_*(*m*_0_, *h*_0_) = *D_m,m_*(*m*_0_, *h*_0_; *m*_0_, *h*_0_), *C_hh_*(*m*_0_, *h*_0_) = *D_h,h_*(*m*_0_, *h*_0_; *m*_0_, *h*_0_), *C_mh_*(*m*_0_, *h*_0_) = *D_m,h_*(*m*_0_, *h*_0_; *m*_0_, *h*_0_) and to the leading order

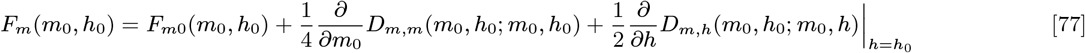

and

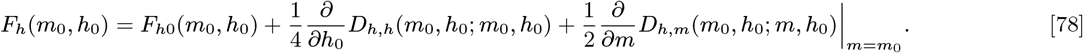

Note that 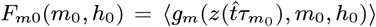 and 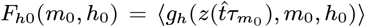. The integral in Eq. (76) to the leading order can be computed as

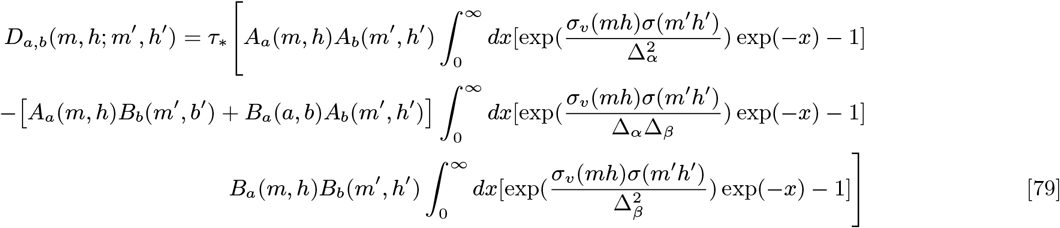

where 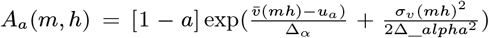, 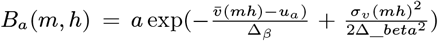 and *τ*_*_ is the effective timescales of noise due to both *m* and *h* fluctuations and it corresponds to

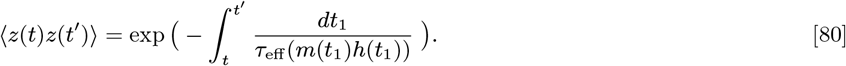

To compute the integral in Eq. (80) to determine *τ*_*_, we assume *t* < *t′* and the slow dynamics of *m* and *h* indicate that *m*(*t*) = *m*,*h*(*t*) = *h*, *m*(*t′*) = *m′* and *h*(*t′*) = *h′*. Thus, to the leading order

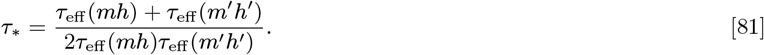

This determines all terms for the two-dimensional Fokker-Planck equation in Eq. (75) to describe the stochastic dynamics of the slow fluctuations in *m* and *h*.

##### One-dimensional Fokker-Planck equation for evolution of slow *h* when *m* is fast

In this regime, we do not expect to see an increase in the timescales of the fluctuation. However, we can still characterize the system via a one-dimensional Fokker-Planck equation for the evolution of slow *h* density. As described in the main text, we need to derive the evolution of *v* density and then use the statistics of rapid fluctuation in *v*; we drive the Fokker-Planck equation for the evolution of slow *h* distribution. Here, we have

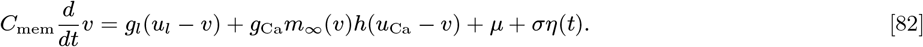

Note that the distribution of *v* cannot be Gaussian due to its coupling with *m*_∞_(*v*), however we can drive P(*v*; *t*) as Gaussian expansions. To do so, it is useful to apply a change of variable 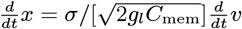. Therefore,

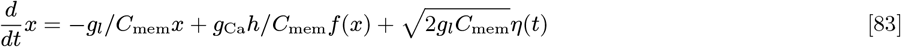

where, 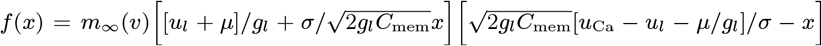 and the equivalent Fokker-Planck equation is

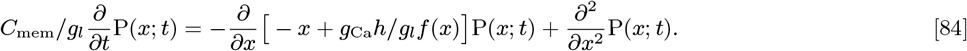

It is straightforward to drive the equilibrium distribution Peq(*x*). The time dependent solution to this equation in terms of Hermite polynomials, He*n*(*x*), can be written as

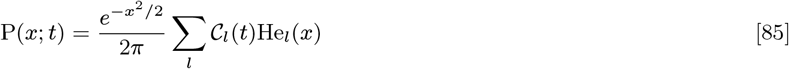

where 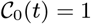 and for *l* ≠ 0 we have

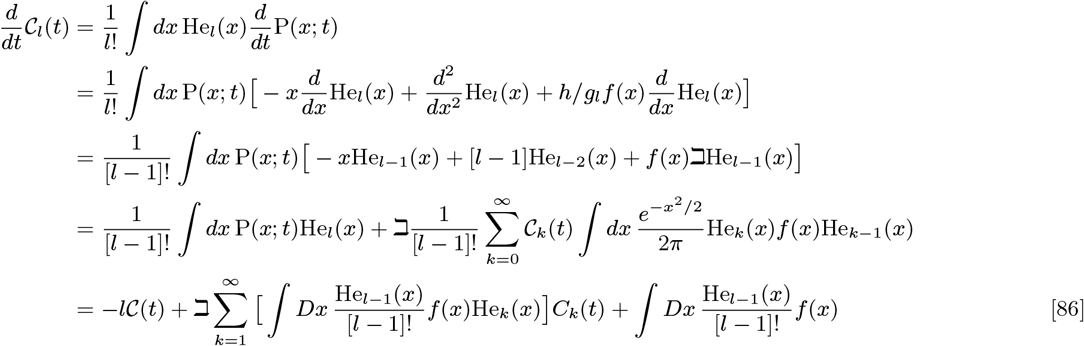

where 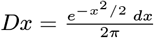 and 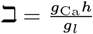. Thus,

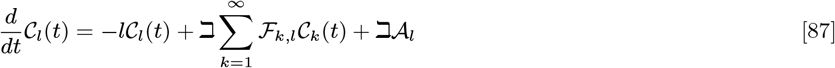

where 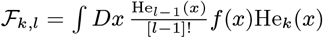 and 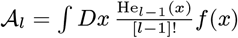. At equilibrium we have

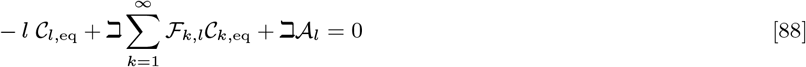

Using matrix notation, we have at equilibrium

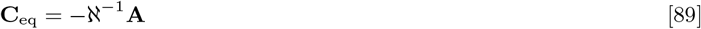

where matrix ℵ = ℶ[D + ℶF]. This allows us to express 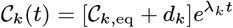, where *d_k_* is the off-set in the dynamics due to the non-zero vector **A** and it corresponds to *k*-th element in the vector of −ℵ^−1^**A** and *λ_k_* is the *k*-th eigenvalue and 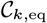 is the corresponding eigenvectors of the matrix ℵ. Assuming initially 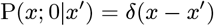, we have

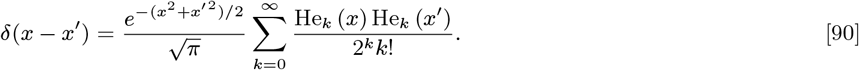

Now, using Eq. (85), we determine

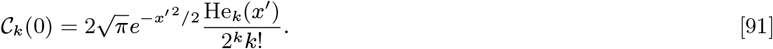

Thus,

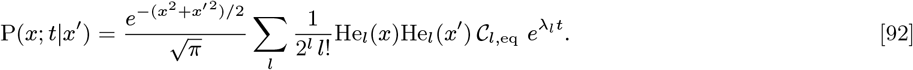

Since the equilibrium voltage distribution is stable (all *λ_l_* ≤ 0), hence the series is converging quickly. The conditional distribution of the re-scaled voltage further allows to determine fluctuation in *h*. On the 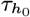 timescale, these fluctuations can be approximated by a white noise process. Therefore, the Langevin equation for *h* in that time scale specifies by

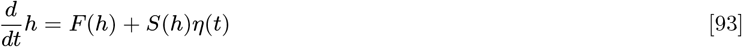

where, 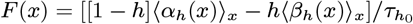 (note that 〈.〉*_x_* indicates the averaging over the equilibrium distribution of *x*, P_eq_(*x*)), and

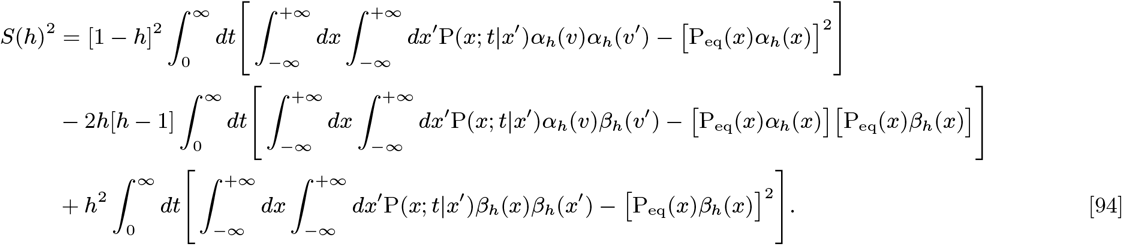

The integration over *t* can be performed analytically,

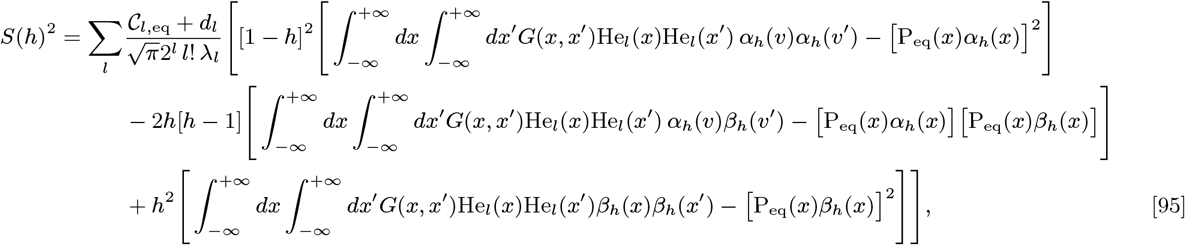

where 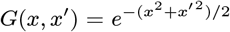. This integral can be numerically evaluated with arbitrary accuracy by the inclusion of *n*-th number of terms in the summation. Note that the prefactor in the summation goes quickly to zero. The corresponding Fokker-Planck equation to the Langevin equation for *h* is

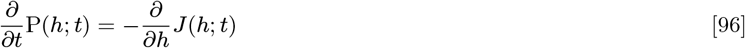

where 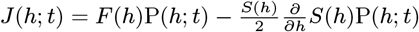 with the boundary conditions of *J* (0; *t*) = 0 and *J* (1; *t*) = 0.

##### Numerical methods for calculation of relaxation time

The Fokker-Planck equations in this work describe the approximate evolution of the probability density of the relevant variable in the system and generally can be written as

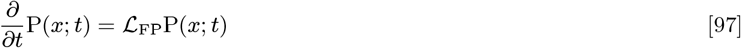

where 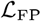 is the Fokker-Planck operator. We use centred finite differences to discretize the operator in the space of variable *x*,

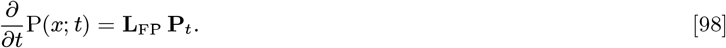

where **P**_*t*_ is the probably vector at time *t* and **L**_FP_ is the Fokker-Planck matrix. Furthermore, note that in our one-dimensional Fokker-Planck description for the activation variable, we require to adjust **L**_FP_ to express boundary conditions ensuring that the probability flux is zero outside of the domain of *m*. In the case of two-dimensional Fokker-Planck equation for evaluation of P(*x, y*; *t*), the discretization can be map also to a two dimensional matrix, **L**_FP_. To do so, we need to concatenate the probability matrix **P**(*i*Δ*x, j*Δ*y*) into vector **P**(*ki,j*), where *ki,j*-th element in the vector corresponds to *i*Δ*x* + *jN* Δ*y*, where *N* is the number of points in the probably grid for each variable. To calculate the eigenvalues of **L**_FP_ we use Arnoldi and Lanczos algorithm implemented in ARPACK.To relaxation time to the equilibrium solution corresponds to the real part of the second largest eigenvalue and defined as 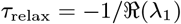.

## ACKNOWLEDGMENTS

FF’s work was supported by the Deutsche Forschungsgemeinschaft (Grant No. FA 1316/2-1). CvW has received funding via CRCNS Grant No. ANR-14-NEUC-0001-01, ANR Grant No. ANR-13-BSV4-0014-02, and No. ANR-09-SYSC-002-01.

